# Elucidation of Puberulic Acid–Induced Nephrotoxicity Using Stem Cell-based Kidney Organoids

**DOI:** 10.1101/2025.04.30.651410

**Authors:** Hiroyuki Nakanoh, Kenji Tsuji, Naruhiko Uchida, Kazuhiko Fukushima, Soichiro Haraguchi, Shinji Kitamura, Jun Wada

## Abstract

**Background and hypothesis:** Recent cases of acute kidney injury (AKI) in Japan have been linked to the Beni-koji CholesteHelp supplement. Renal biopsies from affected patients revealed tubular damage, and puberulic acid was subsequently identified as a potential nephrotoxic contaminant. Recognizing the urgent need for a reliable in vitro nephrotoxicity testing platform, we hypothesized that a kidney organoid-based system could replicate nephrotoxic injury and help identify novel nephrotoxicants. In this study, we developed a screening model using kidney organoids derived from adult rat kidney stem (KS) cells and applied it to evaluate the toxicity of puberulic acid.

**Methods:** Kidney organoids were generated from KS cells and exposed to established nephrotoxicants, including cisplatin and gentamicin, to validate the model. The nephrotoxicity of puberulic acid was evaluated using both KS cell-derived organoids and wild-type mice. Nephrotoxicity was assessed by morphological changes, Kim-1 mRNA expression, transmission electron microscopy (TEM), and analysis of markers related to mitochondrial injury, oxidative stress, and apoptosis.

**Results:** The kidney organoids reproduced morphological features of injury induced by known nephrotoxins and showed significant upregulation of Kim-1 mRNA. Puberulic acid-treated organoids displayed ultrastructural features consistent with acute tubular necrosis (ATN) and increased Kim-1 expression. In vivo, puberulic acid-exposed mice exhibited impaired renal function and histological findings consistent with ATN. Both in vitro and in vivo models revealed mitochondrial structural abnormalities and reduced expression of cytochrome c oxidase subunit IV (COX-IV). Additionally, oxidative stress and apoptotic markers, including 8-hydroxy-2’-deoxyguanosine (8-OHdG) and cleaved caspase-3, were significantly elevated, suggesting that puberulic acid induces mitochondrial dysfunction and oxidative stress, leading to tubular cell death.

**Conclusion:** Puberulic acid-induced nephrotoxicity was demonstrated using our kidney organoid model. KS cell-derived kidney organoids provide a simple, reproducible, and rapid platform for nephrotoxicity assessment, which contribute to a reduction of animal use in toxicology research.

## Introduction

In line with the 3R principles, replacement, reduction, and refinement, organoid-based New Approach Methodologies (NAMs) may serve as alternatives to animal testing in toxicity studies [1]. In the field of nephrotoxicity assessment, kidney organoids derived from induced pluripotent stem (iPS) cells have emerged as a promising in vitro alternative to animal-based models [2] [3]. However, their application in detecting nephrotoxicants remains limited. We have previously established kidney stem (KS) cells derived from a stem/progenitor-like cell line originating from the S3 segment of the renal proximal tubule in adult rats [4]. These KS cells are capable of differentiating into both proximal and distal tubule structures in a three-dimensional culture system [5], offering a suitable model for nephrotoxicity screening.

Recently, acute kidney injury (AKI) associated with *Beni-koji CholesteHelp*, a cholesterol-lowering supplement manufactured by Kobayashi Pharmaceutical and containing red yeast rice (Beni-koji), has emerged as a significant public health concern in Japan [6-10]. Renal biopsies in affected cases revealed tubular injury suggestive of direct nephrotoxicity ^10^. A recent investigation by Japan’s the National Institute of Health Sciences and identified that the implicated product batches contained puberulic acid [11]. Further analysis confirmed that this compound can be synthesized by Penicillium adamezioides, a blue mold detected in the production facility [12]. Therefore, Chemical contamination with puberulic acid has been proposed as a potential cause. Puberulic acid has previously been studied for its antimalarial properties, and earlier reports indicate that it is lethal to mice following two intraperitoneal administrations at 5 mg/kg [13] [14]. However, the nephrotoxic properties and underlying mechanisms of puberulic acid remain poorly understood.

In a prior study, we demonstrated that a specific lot of the *Beni-koji CholesteHelp* supplement induced pathological tubular injury in our KS cell–derived kidney organoid model [15]. In the present study, we refined this model to enable quantitative assessment of tubular damage by measuring *Hepatitis A Virus Cellular Receptor 1* (*HAVCR1*), also known as kidney injury molecule-1 (Kim-1), mRNA expression via real-time PCR. Using this system, we investigated the nephrotoxicity of puberulic acid in vitro and further validated the findings in an in vivo mouse model.

## Materials and Methods

### Cell culture and organoid formation

The KS cells were maintained on type I collagen (IWAKI AGC Techno Glass Corporation, Japan) under previously described undifferentiated culture conditions [4]. The KS cell-derived organoids were generated as previously described [5].

Briefly, cells were trypsinized, harvested, and resuspended in medium, ensuring minimal residual trypsin to avoid interference with cell aggregation. Cell suspensions were seeded onto the plates by hanging drop method, with 2.0×10^5^ cells in each 25-30 µL droplet per well. The clusters were incubated in 5% CO_2_ and 100% humidity at 37°C for 8 hours. For three-dimensional culture, 200 µL of a half-Matrigel solution was applied to Transwell inserts (Corning Life Sciences, USA) and placed into 24-well plates containing 400 µL of differentiation medium. The half-Matrigel solution consisted of a 1:1 mixture of Matrigel and differentiation medium (DMEM/F-12 supplemented with 10% fetal bovine serum (FBS), 250 ng/mL hepatocyte growth factor (HGF), 250 ng/mL glial cell line-derived neurotrophic factor (GDNF), 250 ng/mL basic fibroblast growth factor (bFGF), 250 ng/mL epidermal growth factor (EGF), and 250 ng/mL bone morphogenetic protein-7 (BMP-7). KS cell clusters were transferred into the half-Matrigel solution and cultured at 37°C with 5% CO_2_ and 100% humidity for 2-3 weeks. Cisplatin (60 µM; FUJIFILM Wako Pure Chemical Corporation), gentamicin (10 mM; Selleck Chemicals, USA), and puberulic acid (2, 10 and 50 µM) were applied to the organoids, and were cultured in differentiation medium for 72 hours.

### Animal experiment

The experimental protocol was approved by the Animal Ethics Review Committee of the Okayama University Graduate School of Medicine, Density and Pharmaceutical Sciences (OKU-2024474). Eight-week-old male C57BL/6N mice (CLEA Japan, Japan) were housed with free access to tap water and standard laboratory chow. All animals were cared for in accordance with the relevant guidelines and regulations. Mice were randomly assigned to the following two groups (n = 6 per group): the control group (CTR) and the puberulic acid group (PUB). Mice in the PUB group were intraperitoneally injected with puberulic acid (5.0 mg/kg body weight) on days 0 and 1. Mice in the CTR group were intraperitoneally injected with 100 μL saline on days 0 and 1. All mice were euthanized on day 4, and blood and kidneys samples were harvested for analysis.

### Blood and Urine Biochemical Measurements

Mouse blood samples were collected immediately before euthanizing, and serum creatinine and Blood urea nitrogen (BUN) were measured at the Department of Animal Resources, Advanced Science Research Center, Okayama University. Mouse urine samples were centrifuged to remove cells, cell debris, and other contaminants. Urinary albumin was measured with the Mouse Urinary Albumin Assay Kit (FUJIFILM Wako Pure Chemical Corporation, Japan), and urinary creatinine was measured with a creatinine kit (FUJIFILM Wako Pure Chemical Corporation, Japan).

### Histological and immunofluorescence

Kidney organoids and mouse kidneys were fixed with 10% formalin (Nacalai Tesque, Japan), embedded in paraffin, sectioned at 4 µm, and stained with Hematoxylin and eosin (H&E) or periodic acid-Schiff (PAS). Tubular damage in mouse kidneys was scored by calculation of the percentage of tubules in the corticomedullary junction that displayed tubular dilatation, tubular atrophy, tubular cast formation, and sloughing of tubular epithelial cells or loss of the brush border. The scoring criteria were as follows: 0, none; 1, ≤10%; 2, 11–25%; 3, 26–45%; 4, 46– 75%; and 5, >76%. At least 10 high-power fields (magnification, ×200) per section for each sample were examined (n=6 per group) [16, 17]. For immunofluorescence, deparaffinized sections underwent antigen retrieval with EDTA, blocked with 1% BSA-PBS, and incubated with primary antibodies [fluorescein-labeled lotus tetragonolobus lectin (LTL) (Invitrogen), rabbit cleaved caspase-3 (Cell Signaling Technology, USA), and goat Kim-1 (R&D systems, USA) as previously described [18]. For cytochrome c oxidase subunit IV (COX-IV) staining, after antigen retrieval, sections were fixed with 4% paraformaldehyde (PFA) and subsequently permeabilized with 0.1% Triton X-100. For 8-OHdG staining, sections were similarly fixed with 4% PFA following antigen retrieval, then treated with 2 M hydrochloric acid and neutralized with Tris buffer. 4’,6’-diamidino-2-phenylindole (DAPI) (Roche Diagnostic, Switzerland) counterstain were used to visualize nuclei. Imaging was performed with FSX-100 (Olympus, Japan). Cleaved-caspase 3-positive cells per field were evaluated in three kidney organoids tissue sections using five randomly selected images at x200 magnification.

### Transmission Electron Microscopy (TEM)

TEM was performed as described previously [5]. Briefly, specimens were fixed in 0.1 M cacodylate buffer containing 2.5% glutaraldehyde (pH 7.2) and processed thin sections were stained with uranyl acetate and citrate and examined using a Hitachi H-700 electron microscope.

### Western Blotting

Western blotting analysis was performed as previously described [19]. Briefly, total proteins were extracted using RIPA buffer (Thermo Fisher Scientific, USA) supplemented with protease inhibitors (Promega, USA). Protein concentration was determined using a BCA Protein Assay Kit (Thermo Fisher Scientific, USA). Equal amounts of protein were separated by SDS-PAGE, transferred to nitrocellulose membranes, and blocked with 5% skim milk. Membranes were incubated overnight at 4°C with primary antibodies [GAPDH (Cell Signaling Technology, USA), Kim-1 (R&D systems, USA)], Cox-IV (Proteintech, USA) and 8-hydroxy-2’-deoxyguanosine (8-OHdG) (Japan Institute for the Control of Aging, Japan), followed by HRP-conjugated secondary antibodies. Signal detection was performed with an enhanced chemiluminescence system (GE Healthcare, USA) and imaged using the Amersham Imager 600. Densitometric analysis was performed using ImageJ software.

### Real-time RT-PCR

RNA isolation and qPCR were performed using the kidney organoids (n = 3 per group), as previously described [20]. Total RNA was extracted using an RNA extraction kit (Qiagen Sciences, USA). Real-time PCR was performed on a StepOnePlus™ Real-Time PCR System (Applied Biosystems, USA) using TaqMan Fast Advanced Master Mix (Applied Biosystems, USA). The amplification protocol included an initial enzyme activation at 95 °C for 20 seconds, followed by 40 cycles of denaturation at 95 °C for 1 second and annealing at 60 °C for 20 seconds. The comparative threshold cycle (Ct) method was used to calculate fold amplification. Gene expression was analyzed via qPCR using primers specific for HAVCR1 (assay ID: Rn00597703_m, Thermo Fisher Scientific, USA). Appropriate negative and positive controls were included to validate qPCR performance.

### Cell viability analysis

The immortalized human proximal tubular epithelial cell line HK-2 (American Type Culture Collection, USA) was cultured in DMEM/F12 supplemented with 10% FBS, were seeded in 96-well plates and incubated overnight at a density of 5,000 cells per well in 100 μL of medium to adhere. Then, puberulic acid was added at various concentrations (0.4-50 μM) and incubated for 24 hours. The cell viability following puberulic acid treatment was assessed using Cell counting kit-8, according to the manufacturer’s protocol (Dojindo Laboratories, Japan), as previously described [21]. Absorbance at 450 nm was measured using a microplate reader (Thermo Fisher Scientific, USA), and cell viability was expressed as a percentage relative to the control group (n=8 per group).

### Statistical analysis

All data are presented as means ± standard error of the mean (SEM).

Statistical analysis was performed using JMP software (JMP® 13.2, SAS Institute). Comparisons between two groups were made using either the Student’s t-test or the Wilcoxon rank-sum test, as appropriate. A p-value of < 0.05 was considered as significantly different.

## Results

### Evaluation of Tubular Injury on Kidney Organoids

Kidney organoids were generated using KS cells, and the tubular injury was evaluated following treatment with cisplatin and gentamicin (Figure 1A). Histological evaluation by stereomicroscopy revealed tubules cell detachment, narrowing and structural disruption in the cisplatin-treated group at 72 hours. H&E staining confirmed detachment of tubular epithelial cells, along with cellular and nuclear degeneration in both the cisplatin-and gentamicin-treated groups (Figure 1B). Immunofluorescence staining demonstrated increased expression of Kim-1 in both treatment groups (Figure 1C). These findings were further supported by qPCR analysis, which showed significant upregulation of Kim-1 (*HAVCR1*) mRNA expression compared to the control group (Figure 1D). In addition, immunofluorescence revealed a significant increase in cleaved caspase-3–positive apoptotic cells in the tubules of treated kidney organoids (Figure 1, C and E).

**Figure 1.**
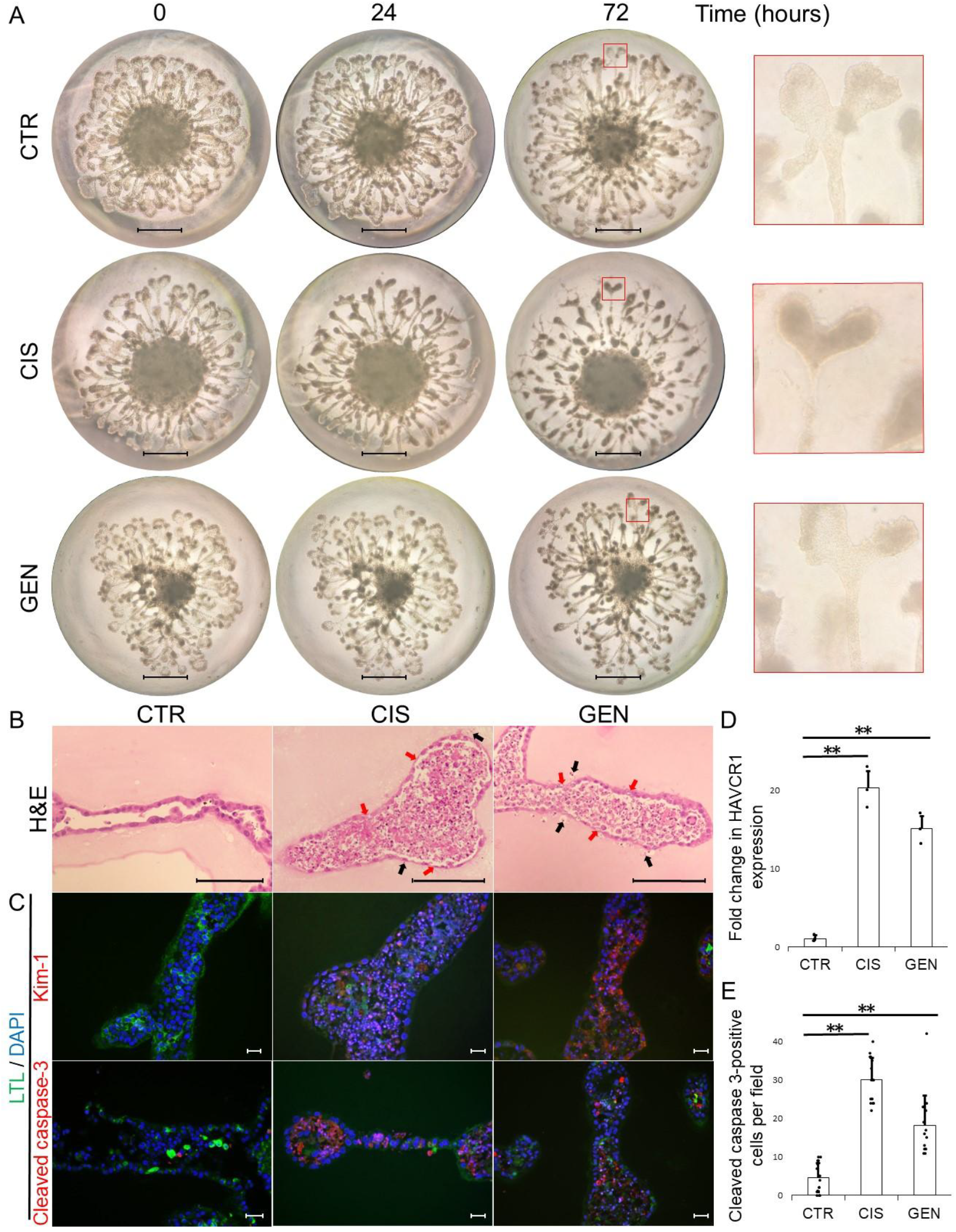
Tubular Injury in Kidney Organoids Induced by Cisplatin and Gentamicin. (A) Time-course morphological changes of kidney organoids treated with cisplatin (CIS) or gentamicin (GEN). Organoids were cultured under control conditions (CTR) or exposed to CIS or GEN, and observed at 0, 24, and 72 hours. Morphological alterations such as tubular cell detachment, narrowing, and disruption became progressively evident, particularly in the CIS group at 72 hours. Enlarged images on the far right highlight structural abnormalities. Scale bar, 1 mm. (B-E) Histological and immunofluorescence analysis of kidney organoids after 72-hour treatment. (B) Hematoxylin and eosin (H&E) staining of kidney organoids. Tubular epithelial cell detachment (black arrows) and degeneration (red arrows) were observed in both the CIS and GEN groups. (C) Immunofluorescence staining for kidney injury molecule-1 (Kim-1, red) and cleaved caspase-3 (red), co-stained with Lotus tetragonolobus lectin (LTL, green) and DAPI (blue). Both Kim-1 and cleaved caspase-3 expression were markedly increased in the CIS and GEN groups compared to CTR. Scale bar, 20 μm. (D) Quantitative real-time PCR analysis showing significant upregulation of Kim-1 (*HAVCR1*) mRNA in the CIS and GEN groups. Expression was normalized to Gapdh and presented as fold change relative to CTR (n = 3 per group). (E) Quantification of cleaved caspase-3–positive cells per field. Five randomly selected fields from three independent samples were analyzed per group. Data are shown as mean ± SEM. Statistical analysis was performed using Student’s *t*-test. ** P < 0.01.

### Puberulic acid induces kidney organoid damage

To evaluate the nephrotoxic potential of puberulic acid, we first assessed the viability of human proximal tubular cells (HK-2 cells), which revealed significant cytotoxicity at a concentration of 50 μM (Supplemental Figure S1). We next utilized the kidney organoid system to further investigate its effects. Treatment with puberulic acid (50 μM) result in tubular detachment, as observed by stereomicroscopy and confirmed by H&E staining (Figure 2A-B). TEM revealed ultrastructural changes, including loss of intercellular junctions and the presence of low-electron-density vesicles containing remnants of organelles within dilated vacuoles, likely representing lysosomes at 10 and 50 μM (Figure 2C). These morphological changes were consistent with findings previously reported in kidney organoids treated with *Beni-koji Cholestehelp* supplement, and resemble features of drug-induced acute tubular necrosis observed in human kidneys. In addition, Kim-1 expression was elevated in the puberulic acid-treated group, as confirmed by immunofluorescence, western blotting, and qPCR (Figure 2, D-F).

**Figure 2.**
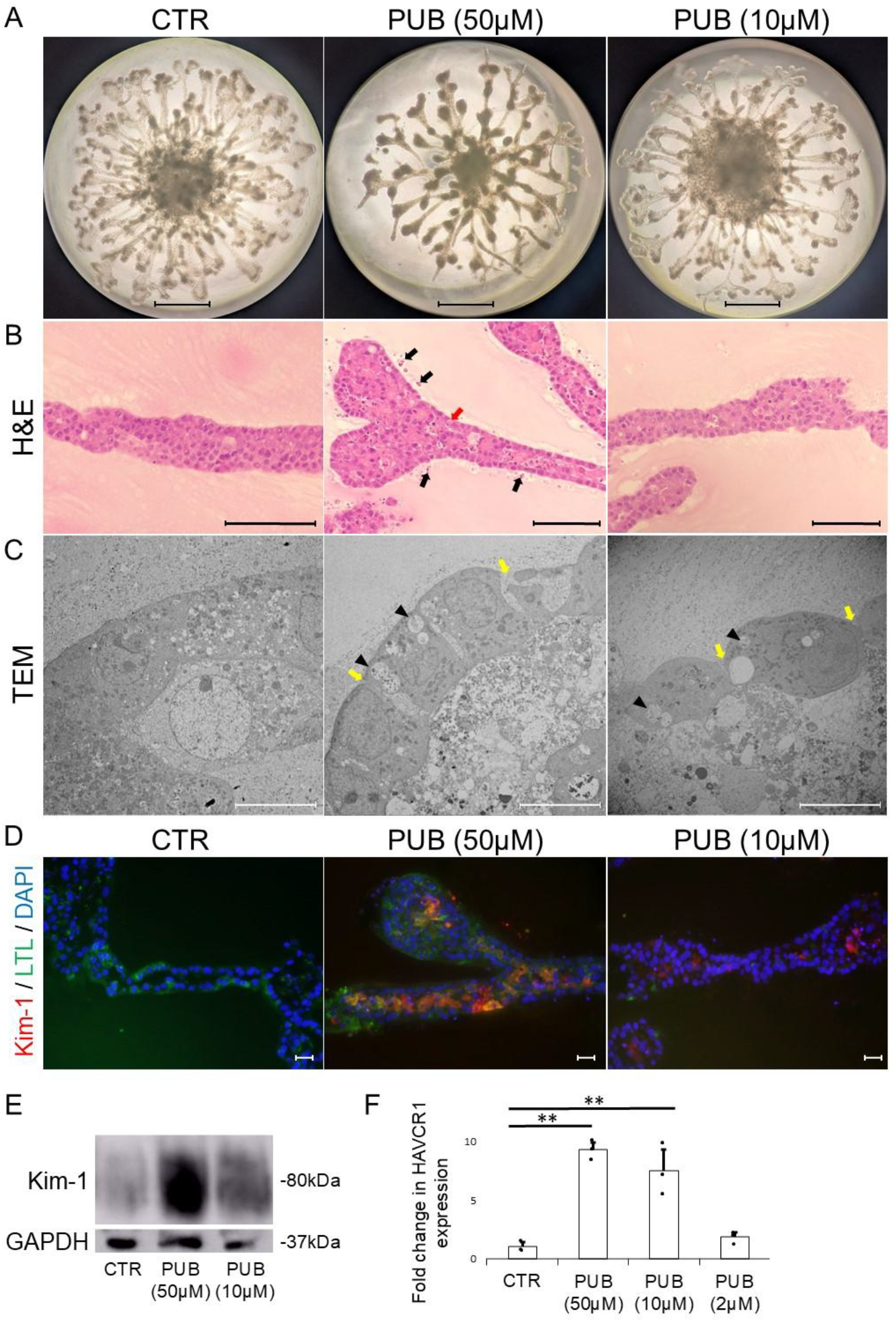
Tubular Injury in Kidney Organoids Induced by Puberulic Acid. (A) Stereomicroscopy images of kidney organoids cultured under control conditions (CTR) or treated with puberulic acid (10 μM and 50 μM) for 72 hours. Tubular cell detachment was observed in the PUB group at 50 μM. Scale bars: 1 mm. (B) Hematoxylin and eosin (H&E) staining showing detachment of tubular epithelial cells (black arrows) and cellular degeneration (red arrows) in the PUB group. Scale bars: 100 μm. (C) Transmission electron microscopy (TEM) images revealing intercellular separation (yellow arrows) and vacuolation with low-electron-density vesicles (arrowheads) in the PUB groups. Scale bars: 10 μm. (D) Immunofluorescence staining for kidney injury molecule-1 (Kim-1, red) and cleaved caspase-3 (red) in Lotus tetragonolobus lectin (LTL, green)-positive tubules. Nuclei were counterstained with DAPI (blue). Kidney organoids were untreated (CTR) or treated with puberulic acid (10 μM and 50 μM) for 72 hours. Increased Kim-1 expression was observed in PUB-treated groups compared to the CTR group. Scale bars: 20 μm. (E) Western blot analysis showing upregulation of Kim-1 protein expression in PUB-treated kidney organoids compared to CTR. (F) Quantitative real-time PCR analysis showing significantly increased Kim-1 (*HAVCR1*) mRNA expression in PUB-treated organoids (2, 10, and 50 μM) compared to CTR. Gene expression was normalized to Gapdh and expressed as fold change relative to the CTR group. Data are presented as mean ± SEM (n = 3 per group). Statistical analysis was performed using Student’s *t*-test. ** P < 0.01.

### Puberulic acid induces acute kidney injury in mice

To validate the nephrotoxic effects of puberulic acid in vivo, we administered the compound to C57BL/6N mice (5.0 mg/kg, on days 0 and 1). Mice treated with puberulic acid (5.0 mg/kg, days 0 and 1) showed elevated levels of serum creatinine and urinary albumin-to-creatinine ratios (Figure 3A, 3C). PAS staining revealed tubular atrophy, shedding, and cast formation, suggestive of acute tubular necrosis, while no glomerular abnormalities were evident (Figure 3D). The renal tubular injury score was significantly higher in the PUB group compared to the CTR group (Figure 3E). TEM further demonstrated vacuolar degeneration of proximal tubular cells, loss of brush borders, and mitochondrial degeneration in the PUB group, consistent with tubular epithelial cell necrosis (Figure 3D). In contrast, no ultrastructural abnormalities were detected in the glomeruli, distal tubules, or collecting ducts in either the PUB or CTR groups (Supplemental Figure S2). Moreover, immunofluorescence staining revealed marked upregulation of Kim-1 in the proximal tubules of the PUB group (Figure 3F), which was further confirmed by western blot analysis (Figure 3G).

**Figure 3.**
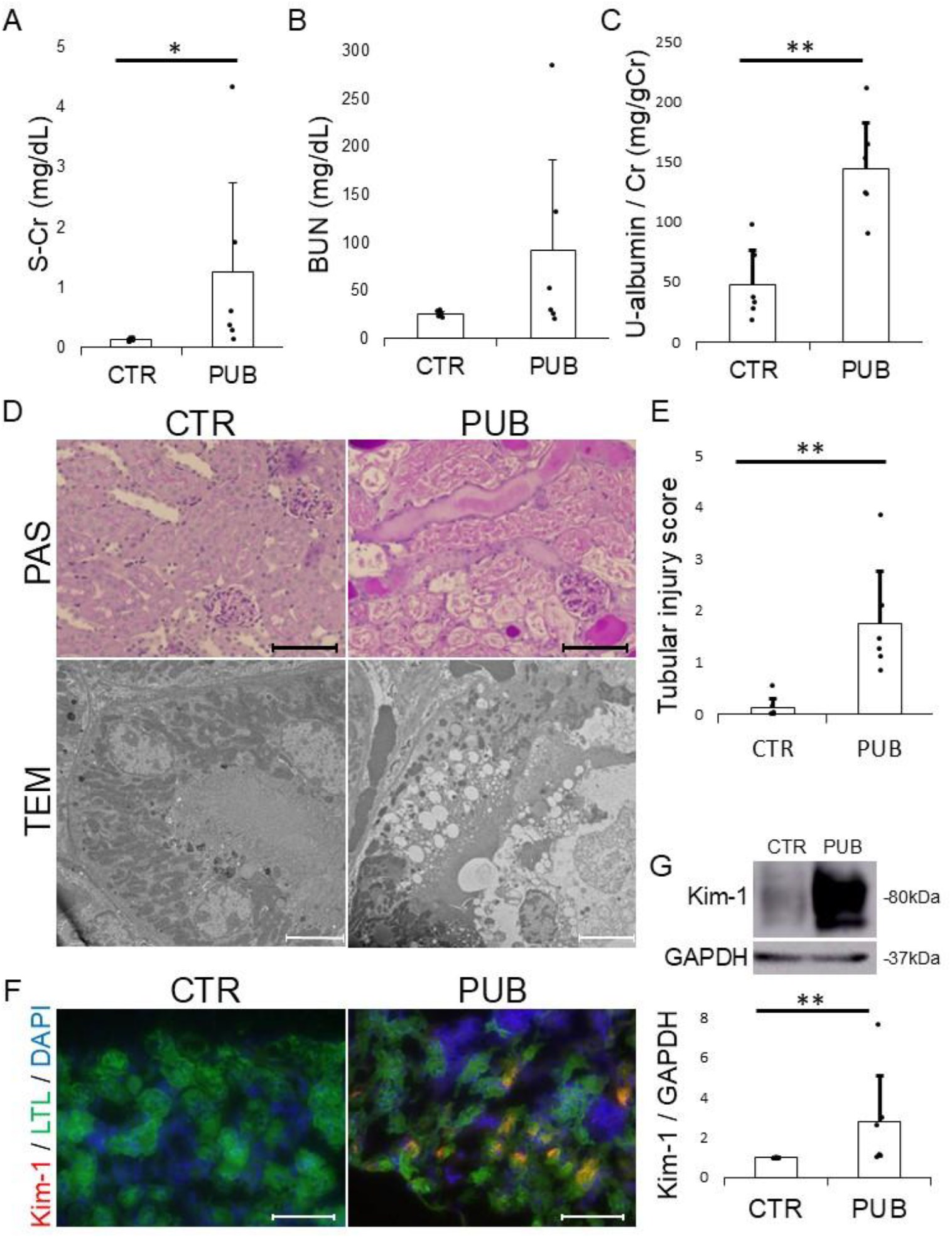
Puberulic Acid Induces Acute Kidney Injury in Mice. Histological and functional evaluation in mice under control conditions (CTR) or treated with puberulic acid (PUB) for 4 days (n = 6 per group). (A) Serum creatinine (S-Cr) levels were significantly elevated in the PUB group compared to the CTR group. (B) Blood urea nitrogen (BUN) levels were higher in the PUB group, although the difference was not statistically significant. (C) The urinary albumin-to-creatinine ratio (U-albumin/Cr) was significantly increased in the PUB group compared to the CTR group. (D) Upper panels: Periodic acid– Schiff (PAS) staining. Scale bars: 5 μm ; lower panels: transmission electron microscopy (TEM) images showing features consistent with acute tubular necrosis in the PUB group. Scale bars: 100 μm. (E) Tubular injury scores were significantly higher in the PUB group than in the CTR group. (F) Immunofluorescence staining of kidney sections showing increased expression of kidney injury molecule-1 (Kim-1, red) in the PUB group compared to the CTR group. Nuclei were counterstained with DAPI (blue). Scale bars: 100 μm. (G) Western blot analysis demonstrating upregulation of Kim-1 protein in the PUB group. Data are presented as mean ± SEM. Statistical analysis was performed using Student’s t-test or Wilcoxon rank-sum test. * P < 0.05, ** P < 0.01.

### Puberulic acid induces tubular mitochondrial dysfunction

TEM revealed distinct ultrastructural alterations in tubular epithelial cells following puberulic acid treatment. In the CTR group, both mouse kidney and kidney organoid samples exhibited well-preserved mitochondria with intact cristae and an organized cellular architecture. In contrast, samples from the puberulic acid-treated (PUB) group exhibited a notable reduction in mitochondrial number, accompanied by structural abnormalities such as swelling and loss of cristae (Figure 4A). These findings indicate mitochondrial damage and are consistent with early stage of tubular epithelial cell apoptosis or necrosis. COX-IV immunostaining demonstrated a decrease in mitochondrial signal intensity within the proximal tubules of both mouse kidneys and kidney organoids in the PUB group compared to the CTR group (Figure 4B). This reduction was further validated by western blot analysis of mouse kidney tissue, which showed significantly decreased COX-IV protein levels in the PUB group (Figure 4C). To evaluate oxidative stress and apoptosis, immunofluorescence analysis was performed. The PUB group showed increased accumulation of 8-OHdG, a marker of oxidative DNA damage, along with a higher number of cleaved caspase-3–positive cells in both mouse kidney and kidney organoid samples (Figures 4B and 4D). Collectively, these findings suggest that puberulic acid induces mitochondrial injury, which subsequently triggers oxidative stress and leads to tubular epithelial cell death.

**Figure 4.**
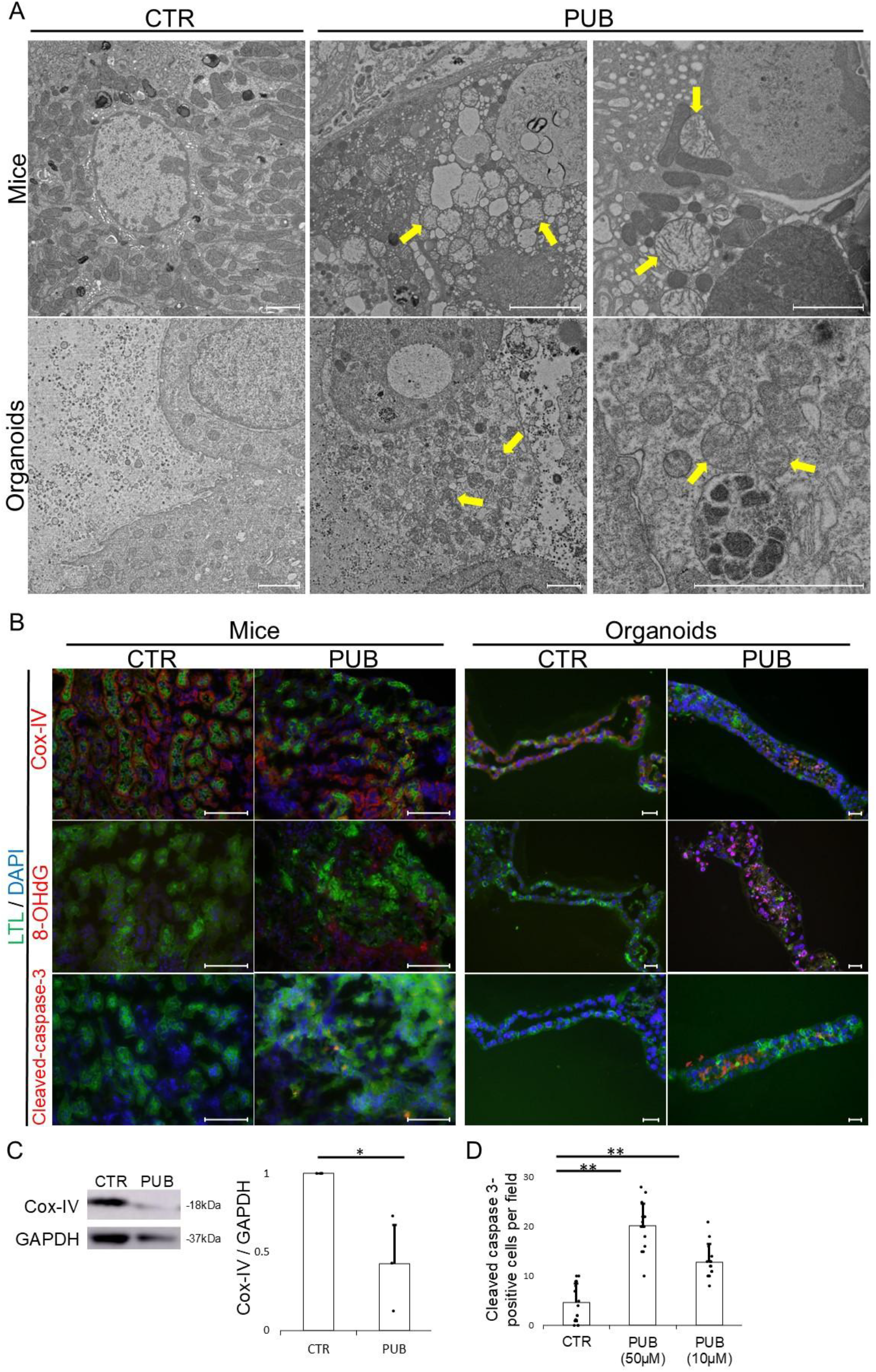
Puberulic Acid Induces Mitochondrial Degeneration. (A) Representative transmission electron microscopy (TEM) images of proximal tubular cells from mouse kidneys (upper panels) and kidney organoids (lower panels) in the control (CTR) and puberulic acid–treated (PUB) groups. Organoids treated with puberulic acid (25 µM). In the CTR group, mitochondria were intact with well-defined cristae and organized structure. In contrast, the PUB group showed reduced mitochondrial number and morphological abnormalities such as swelling along with cristae disruption (yellow arrow). Scale bars: 2 μm. (B) Representative immunofluorescence images of mouse kidney and kidney organoid stained for cytochrome c oxidase subunit IV (COX-IV, red), 8-hydroxy-2’-deoxyguanosine (8-OHdG, magenta), and cleaved caspase-3 (red) in the CTR and PUB groups. Organoids treated with puberulic acid (50 µM). Lotus tetragonolobus lectin (LTL, green) marks proximal tubules, and nuclei were counterstained with DAPI (blue). Compared to the CTR, the PUB group showed a marked reduction in COX-IV signal intensity, increased accumulation of 8-OHdG, and elevated cleaved caspase-3–positive cells. Scale bars: 100 μm (mice), 20 μm (organoids). (C) Western blot analysis of kidney tissue lysates showing decreased COX-IV protein levels in the PUB group compared to the CTR group. GAPDH was used as a loading control. Quantitative densitometry is shown as mean ± SEM (n = 3). Statistical analysis was performed using Student’s *t*-test. *P < 0.05. (D) Quantification of cleaved caspase-3–positive cells per field. Data represent mean ± SEM (n = 3 organoids per group, five fields each). Statistical analysis was performed using Student’s t-test. **P < 0.01.

## Discussion

We hypothesized that kidney organoids derived from KS cells could serve as a reliable platform to assess nephrotoxicity, particularly for emerging natural toxins such as puberulic acid. In this study, we demonstrated that our organoid model recapitulated tubular injury induced by known nephrotoxins and enabled quantitative evaluation of injury via Kim-1 (*HAVCR1*) expression. Furthermore, puberulic acid induced morphological and molecular features consistent with ATN both in vitro and in vivo. These findings suggest that this organoid model is suitable for screening nephrotoxicity and investigating toxic mechanisms.

KS cell-derived kidney organoids were previously developed as a screening tool for nephrotoxicity based on morphological assessment ^[15, 22]^. To enhance the reproducibility of nephrotoxicity screening, we introduced qPCR-based quantification of Kim-1 (*HAVCR1*) expression. This enhancement adds reproducibility and objectivity to the assessment. Previous studies using iPS cell-derived kidney organoids have similarly reported Kim-1 upregulation following gentamicin exposure [23]. Recent advances, including microwell-based automated culture systems, have further expanded the utility of kidney organoids in high-throughput screening for nephrotoxicity and nephroprotection [2] [3]. Compared to iPS cell-derived systems, the KS cell-derived model offers a rapid, reproducible, and cost-effective alternative. In our previous report, we demonstrated that specific lots of *Beni-koji CholesteHelp* supplements induced renal tubular injury in this organoid model ^15^. However, its nephrotoxicity remained poorly understood. In this study, we demonstrated that puberulic acid induces nephrotoxicity, supporting its role as a potential causative agent of AKI associated with specific lot of *Beni-koji CholesteHelp* supplement.

Mitochondria generate the energy compound ATP via oxidative phosphorylation, and mitochondrial dysfunction results in excessive production of proinflammatory and injurious molecules, such as reactive oxygen species (ROS), leading to oxidative stress that activates both necrotic and apoptotic cell death pathways [24] [25]. This pathogenic cascade has been documented in AKI, including cisplatin-induced nephrotoxicity, which is reported to be associated with a decrease of cytochrome c oxidase (COX) activity and a reduction in COX-IV protein expression, ultimately resulting in respiratory chain dysfunction and increased mitochondrial ROS production [25]. In the present study, mice treated with puberulic acid exhibited acute proximal tubular necrosis and marked mitochondrial structural abnormalities, as demonstrated by PAS staining and TEM. In addition, decreased expression of COX-IV and elevated levels of 8-OHdG were observed. These findings suggest that puberulic acid induces nephrotoxicity through a pathogenic mechanism similar to that of cisplatin, involving respiratory chain dysfunction and increased mitochondrial ROS production. The kidney organoids closely recapitulated the tubular injury and mitochondrial abnormalities observed in the mouse model, suggesting that they may serve as an alternative to animal experiments in assessing the nephrotoxicity.

Despite its advantages, this kidney organoid system has several limitations. First, these kidney organoids lack the expression of several transporters, including OCT1, OCT2, OAT1, and OAT3 [15, 22]. Consequently, compounds may enter cells via passive diffusion, affecting nephrotoxicity profiles. Second, the absence of immune cells, blood vessels, and endothelial components limits the ability to assess kidney damage related to these factors in the kidney, such as tubulointerstitial nephritis, vascular injury, endothelial damage, and ischemia. Third, since toxins are applied externally rather than through physiological routes such as blood circulation and glomerular filtration, direct extrapolation of drug concentrations to clinical settings is not feasible. Incorporating microfluidic systems or perfusable vasculature may help address this gap and improve translational extrapolation of compound concentrations. Despite these limitations, the kidney organoid model remains a valuable *in vitro* tool for detecting direct tubular toxicity in a controlled environment. It offers a reproducible, scalable, and ethically favorable alternative to animal models, and can be particularly useful in the early-phase nephrotoxicity screening.

In conclusion, this KS cell–derived kidney organoid model represents a suitable and versatile in vitro platform for nephrotoxicity testing and mechanistic studies. The model successfully recapitulates key morphological and molecular features of tubular injury observed in vivo, including those induced by established nephrotoxins such as cisplatin and gentamicin, as well as the naturally occurring compound puberulic acid. These findings validate its utility as a practical and reproducible alternative to animal-based models in preclinical nephrotoxicology. Importantly, the ability to evaluate compound-induced mitochondrial dysfunction, oxidative stress, and cell death pathways in a controlled and scalable system highlights the model’s potential for both hazard identification and mechanistic toxicology. Moreover, this platform aligns with the 3R principles by reducing reliance on animal testing, thereby addressing both ethical and regulatory demands for more human-relevant toxicity assessment systems. Future research should focus on enhancing the physiological relevance of the model, for example by incorporating vascular, immune, and transporter components. In addition, integration with high-throughput and microfluidic technologies may further broaden its utility in drug development and toxicology. With further development, this model may serve as a valuable tool for both predicting human nephrotoxicity and informing personalized and regulatory approaches in toxicology.

## Data availability statement

All data generated or analyzed during this study are included in this published article and its supplementary information files.

## Acknowledgements

None.

## Funding

This work was supported by a Grant-in-Aid for Scientific Research (C) from the Japan Society for the Promotion of Science (JSPS) (grant no. 24K11411, KT) and Wesco Scientific Promotion Foundation (KT). The funder has no influence on the design or analyses in this study.

## Authors’ contributions

H. Nakanoh and K. Tsuji designed the study and drafted the paper; H. Nakanoh, K. Tsuji, N. Uchida, K Fukushima, and S. Haraguchi conducted the experiments; H. Nakanoh and K. Tsuji analyzed the data and made the figures; S. Kitamura and J. Wada reviewed statistical methods and analysis; all authors revised and approved the final version of the manuscript.

## Conflict of interest statement

None declared.

## Notes

### Competing Interest Statement

The authors have declared no competing interest.

### Summary of Updates

Additional analysis of mechanism of nephrotoxicity in kidney organoids.

