## Supplemental Materials for "Elucidation of Puberulic Acid–Induced Nephrotoxicity Using Stem Cell-based Kidney Organoids"

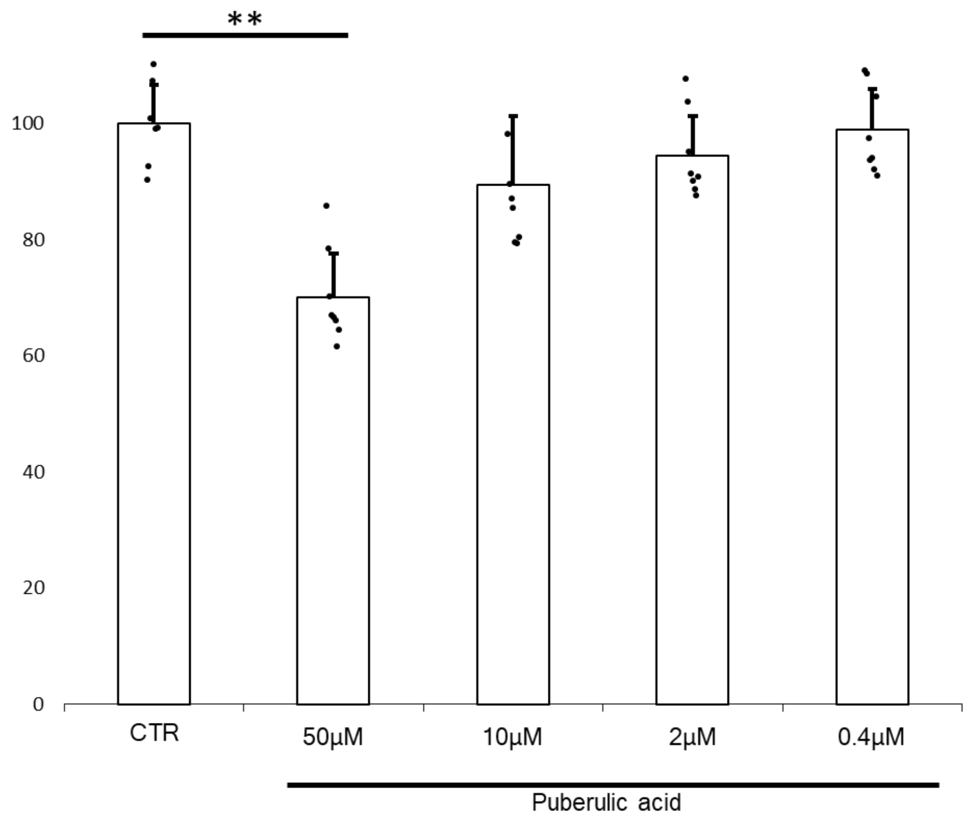


**Supplemental Figure 1.**

A cytotoxicity assay was performed on HK-2 cells treated with puberulic acid at concentrations of 0.4, 2, 10, 50 μM. Cell viability was significantly decreased at 50 μM relative to the control group (CTR) (n=7-8 / group). Values are mean±SD. Analysis was conducted by Turkey test. ** $p<0.01$.

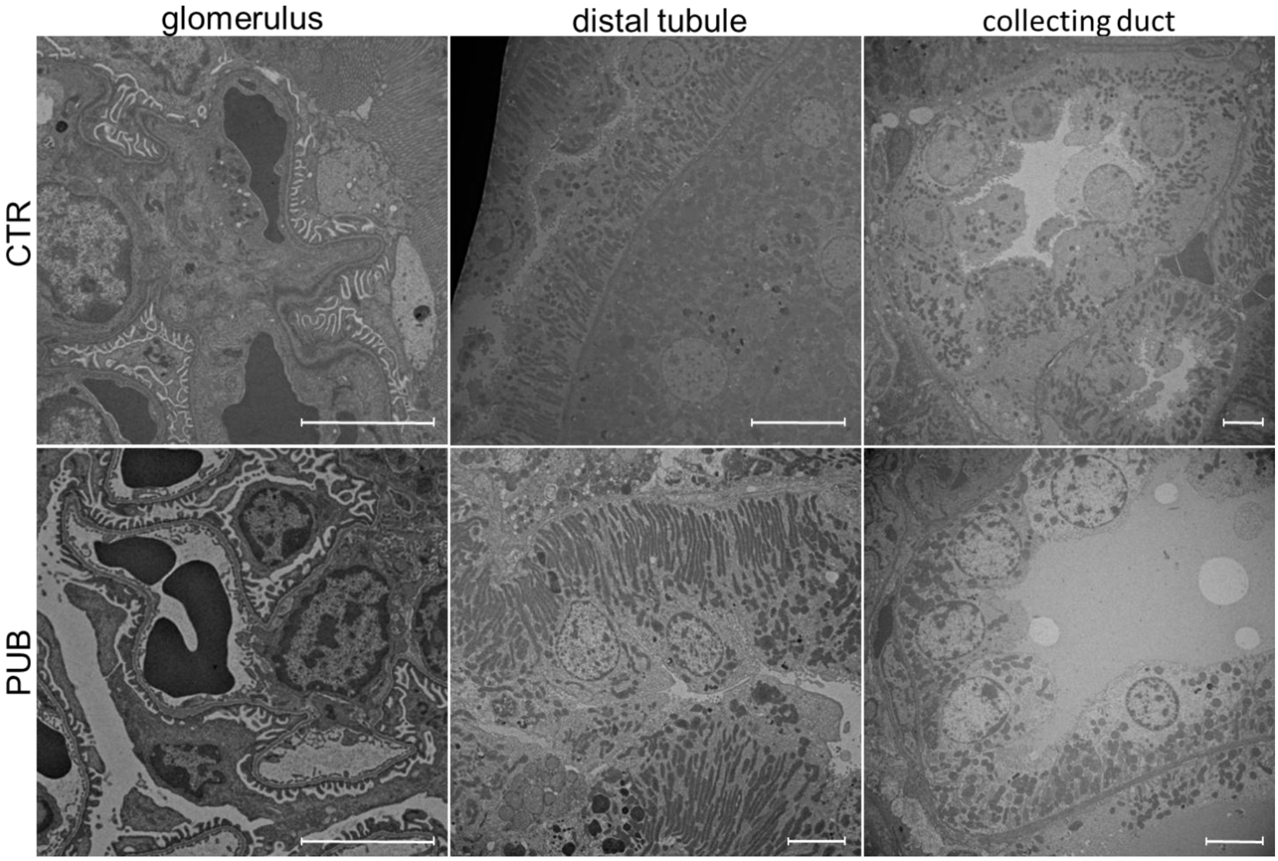


**Supplemental Figure 2.**

TEM images of glomerulus, distal tubule, collecting duct under control conditions (CTR) and treated with puberulic acid (PUB) showing no ultrastructural abnormalities. Scale bars: 5 µm.
